# Swarm-seq Reveals Functional Divergence in CsrA Homologs Across Bacteria

**DOI:** 10.1101/2025.05.05.652322

**Authors:** Jared T. Winkelman, Ethan Yarberry, Jonathan D. Winkelman, Sampriti Mukherjee

**Author notes:** These authors contributed equally to this work (Jared T. Winkelman, Ethan Yarberry).

## Abstract

Bacteria employ sophisticated post-transcriptional regulatory mechanisms to adapt to environmental changes. Carbon storage regulator A (CsrA), a highly conserved small RNA-binding protein, serves as a critical post-transcriptional regulator by recognizing GGA motifs in target transcripts and typically repressing translation. Despite extensive research, how this conserved regulator has evolved diverse species-specific regulatory networks remains unclear. Here, using *Bacillus subtilis* as an experimental chassis, we have developed Swarm-seq — a high-throughput functional genomics platform capable of assessing CsrA homologs across the Bacterial domain for their ability to regulate flagella-dependent swarming motility. Systematically testing over five-hundred codon-optimized *csrA* homologs for flagellin (Hag) repression revealed striking functional divergence, i.e., broadly two classes of CsrAs. Class I, exemplified by CsrAHp from *Helicobacter pylori* and RsmN from *Pseudomonas aeruginosa* strongly inhibited swarming, while Class II, exemplified by CsrAEc from *Escherichia coli* and RsmA from *P. aeruginosa* failed despite high sequence conservation. This differential activity occurred despite all proteins targeting identical GGA motifs in the *hag* transcript’s 5’UTR, indicating evolutionary plasticity in RNA-binding specificity beyond motif recognition. We propose that this plasticity enables a conserved global regulator to develop species-specific regulons through subtle structural adaptations, facilitating diverse physiological responses across bacterial taxa. Our findings establish *B. subtilis* as a powerful platform for characterizing CsrA homologs from genetically intractable bacteria, providing insights into post-transcriptional regulatory network evolution.

## INTRODUCTION

Development of sequencing and computational technologies have led to an abundance of accessible bacterial genomes and promoted advancements in our understanding of gene content in diverse species. However, attributing gene function and regulatory networks across the bacterial domain is limited by our reliance on extrapolation of experimental studies performed in a few model species. Because regulation of gene expression patterns drive adaptation to niches as well as lifestyle decisions such as development of multicellular communities called biofilms (Flemming et al., 2016; Penesyan et al., 2021; Sauer et al., 2022), establishment of pathogenesis (O’Boyle et al., 2020), and antibiotic resistance (Ghosh et al., 2020; Mediati et al., 2021; Palmer et al., 2018), it is imperative to exploit genome mining and develop model-system-free platforms to decipher the mechanisms of conserved global regulators from genome sequences.

Bacteria survive in constantly changing environments and therefore must be able to rapidly alter their gene expression patterns and adjust their proteomes in response to environmental stimuli. Accordingly, post-transcriptional regulation serves as an essential layer to control gene expression tightly and quickly in bacteria and typically involves modulation of messenger RNA (mRNA) elongation, stability, and translation via different types of regulators such as small RNAs (sRNAs) and RBPs (Martínez and Vadyvaloo, 2014). Among RBPs encoded by bacterial genomes, ribonucleases (RNases) assist in the maturation and degradation of mRNAs, while RNA chaperones, RNA helicases, and RNA methyltransferases play key roles in modification of translation efficiency (Holmqvist and Vogel, 2018; Van Assche et al., 2015). One of the most extensively studied bacterial RBPs is CsrA/RsmA (carbon storage regulator A/ regulator of small molecules A), a small (∼60 to 100 amino acids) protein that is widely distributed in bacteria (Fig. 1A). Absence of CsrA often has pleiotropic effects as CsrA serves as a global regulator targeting hundreds of transcripts in various processes such as central carbon metabolism, motility, quorum sensing, the production of extracellular products and biofilm formation (Chatterjee et al., 1995; Lenz et al., 2005; Mukherjee et al., 2011; Sabnis et al., 1995). CsrA is also known to be critical for the regulated expression of virulence factors in pathogens of both animals and plants (Barnard et al., 2004; Gu et al., 2018; Heroven et al., 2008; Vakulskas et al., 2015).

**Fig. 1:**
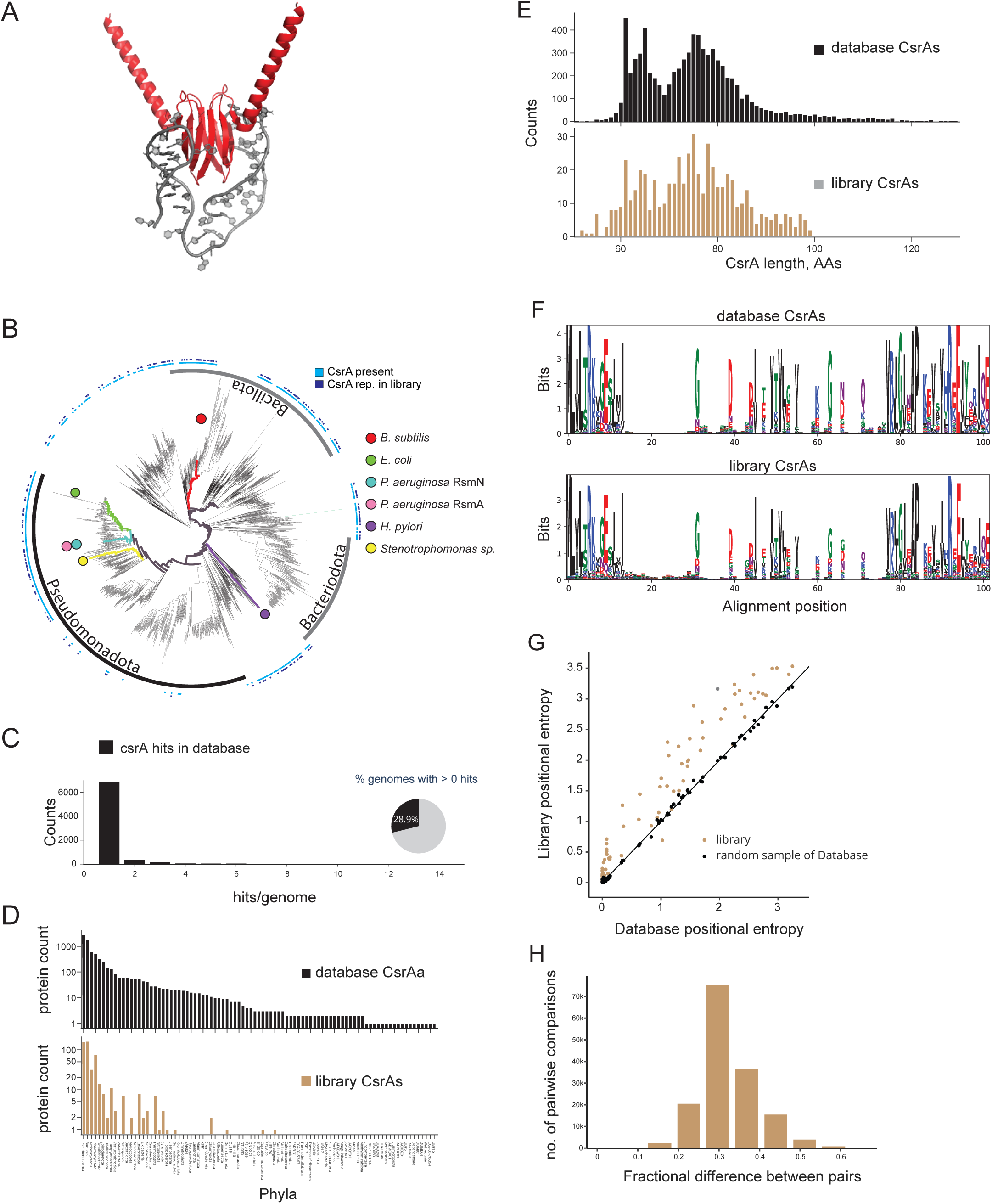
A library of natural CsrAs captures broad structural, phylogenetic, and sequence diversity. **(A)** Structural model *Bacillus subtilis* CsrA bound to an RNA. A predicted structure of *B. subtilis* CsrA (AlphaFold3) is shown interacting with a representative RNA hairpin structure (from PDB 2MF0). This illustrates the conserved mode of CsrA binding to RNA via two hairpins. **(B)** Phylogenetic distribution of CsrAs across bacterial genera. Each leaf represents a genus. Light blue bars indicate where the representative species encodes a CsrA, while dark blue bars show genera with a CsrA included in the synthetic library. Colored circles and branches highlight genera containing CsrAs discussed in more detail in the text. This panel demonstrates the broad taxonomic representation of CsrAs, and the substantial coverage achieved by the library. **(C)** Number of CsrAs per species. Bar plot showing how many distinct CsrAs were identified across species in our full dataset. The inset pie chart shows the proportion of genomes that encode at least one CsrA, indicating its widespread presence across bacteria. **(D)** Phylum-level representation of CsrAs in the full database and the synthetic library. Bar charts show the number of unique CsrAs identified per phylum in the full dataset (top) and the subset selected for the synthetic library (bottom), illustrating how well the library captures phylogenetic breadth. **(E)** Distribution of CsrA protein lengths. Histograms show the distribution of amino acid sequence lengths among CsrAs in the full dataset (top) and the library (bottom). The library reflects the natural size diversity of CsrAs. **(F)** Sequence conservation across CsrAs. Sequence logos generated from all CsrAs in the database (top) and those in our library (bottom) highlight conserved amino acid motifs. The library retains the key residues known to be involved in RNA binding and dimerization. **(G)** Per-position sequence entropy comparison. Positional entropy (a measure of variability at each residue) is plotted for a random sample of full-database CsrAs (black) and the library (tan), matched in size. The increase in the entropy profile indicates that the library contains even more sequence diversity than a random sample. **(H)** Pairwise sequence divergence among library CsrAs. Histogram shows the distribution of pairwise amino acid differences between library CsrAs, demonstrating that the library contains a wide range of divergent sequences.

The canonical mechanism whereby CsrA inhibits translation initiation involves recognition of GGA triplets in the 5’ UTR of its target mRNA to obstruct binding of the 30S ribosomal subunit. In general, the 5′-UTR of the CsrA-regulated mRNAs often contain a high affinity GGA motif that directly overlaps with the ribosome binding site (RBS) and prevents ribosome access (Dubey et al., 2003; Mukherjee et al., 2011). The activity of CsrA is often antagonized by untranslated RNAs such as CsrB and CsrC, which contain multiple binding sites (18 and 9, respectively) that permit sequestration of CsrA (Liu et al., 1997; Weilbacher et al., 2003). The structures of CsrA of *E. coli*, *B. subtilis,* RsmA (regulator of secondary metabolism) of *P. aeruginosa*, RsmE of *P. fluorescens,* and RsmA of *Yersinia enterocolitica* have been determined and present the same overall topology (Gutiérrez et al., 2005; Heeb et al., 2006). The CsrA proteins form a novel class of RNA-binding protein where homodimers consist of two interdigitating monomers (each approx. 61 amino acids, 7KDa). Each RNA binding site is essentially constituted by the first β-strand of one monomer and the fifth β-strand of the other monomer (Mercante et al., 2006).

The aforementioned attributes – small size, broad distribution across genomes, and deep mechanistic understanding in model bacteria – make CsrA amenable for developing a cross-species platform to define molecular determinants of target recognition of widely conserved global regulators in bacteria. Conversely, although heavily studied in the model bacterium *E. coli* (Romeo, 1998; Romeo et al., 1993) and in pathogenic Gamma-proteobacteria such as *P. aeruginosa* (Brencic and Lory, 2009; Chihara et al., 2021; Schulmeyer et al., 2016a)*, Acinetobacter baumannii* (Harding et al., 2018; Singh et al., 2025), and *Vibrio cholerae* (Butz et al., 2021; Lenz et al., 2005), the contribution of this conserved RNA-binding regulator to global gene expression patterns in bacteria from distant phyla remains unclear – a knowledge gap that we strive to address here.

It has been previously hypothesized that regulation of gene expression by CsrA in other species (*i.e. Pseudomonas*, *Erwinia*, *Salmonella*, and *Vibrio*) occurs in a similar manner as in *E. coli* and these CsrAs can complement for each other (Altier et al., 2000; Fortune et al., 2006; Heurlier et al., 2004; Holmqvist and Vogel, 2018). This idea heavily relied on a study that reported complementation of *Pseudomonas fluorescens rsmA* mutant by *Sinorhizobium meliloti csrA* (Agaras et al., 2013). However, recent reports have also highlighted the absence of complementation between CsrAs of closely related species (Kaleta et al., 2022). In sum, the relative dearth of experimental data about post-transcriptional regulatory circuitry across bacteria has hampered the predictive value of known networks.

Here, we present a framework for broad genome scale analysis using CsrA as a candidate global regulator. We combined deep phylogenomic analysis with massively parallel physiological screening in a method we named Swarm-seq - a high-throughput functional genomics platform using *Bacillus subtilis* as a chassis that enables investigation of CsrA homologs from over 500,000 bacterial species for their ability to inhibit flagella-dependent motility. Using Swarm-seq, we uncovered unexpected functional divergence among these phylogenetically related regulators. For instance, CsrAHp from *Helicobacter pylori* and RsmN from *Pseudomonas aeruginosa* potently inhibited swarming, while well-studied homologs CsrAEc from *Escherichia coli* and RsmA from *P. aeruginosa* failed despite high sequence conservation. This differential activity occurred even though all proteins targeted identical GGA motifs in the *hag* transcript’s 5’UTR, revealing evolutionary plasticity in RNA-binding specificity beyond simple motif recognition. Furthermore, our work establishes *B. subtilis* as a robust platform for studying CsrA homologs from genetically intractable bacteria and provides crucial insights into the molecular evolution of post-transcriptional regulatory networks.

## RESULTS

### Development of a phylogenetically diverse library of CsrAs

To build a library that samples as much CsrA diversity as possible, we first sought to understand CsrA primary sequence diversity. CsrA is present in many bacterial genomes, but the extent of its distribution and sequence variation remained unclear. To assess its natural taxonomic and sequence diversity, we first examined CsrA distribution across bacterial taxa. We leveraged the proGenomes database, which contains ∼20,000 curated, nonredundant bacterial proteomes (Mende et al., 2019). Using stringent Hidden Markov Model (HMM) matching criteria, we identified ∼7,500 CsrA homologs from this dataset (Supplementary Fig. 1A). These representatives were found across a wide range of bacterial phyla, including some of the most distantly related branches of the bacterial tree (Fig. 1B, Supplementary Fig. 1B). Approximately 30% of bacterial genomes contained at least one *csrA* gene, with most encoding a single ortholog, although some genomes carried up to 15 (Fig. 1C). While CsrA is broadly distributed and present in most phyla, its prevalence was not uniform across the bacterial tree (Fig. 1D, Supplementary Fig. 1B). It was widely found in certain phyla, such as Bacillota, Planctomycetota, and Pseudomonadota, whereas others, such as Bacteroidota, contained relatively few representatives. These findings highlight both the evolutionary conservation of CsrA and its uneven phylogenetic distribution, suggesting that its regulatory role may be more essential in some lineages than in others.

To systematically investigate the natural diversity of CsrA function, we assembled a library of 555 CsrA homologs from diverse bacterial lineages using UniRef50 clustering, which groups sequences at ≥50% identity to reduce redundancy while maintaining phylogenetic diversity (Table S1). The 555 CsrA representatives in our library span a broad phylogenetic range, demonstrating a diverse representation of CsrA across taxa (Fig. 1B, Supplementary Fig. 1B). Notably, our library does not include CsrA from the *Bacteroidota* and only a few examples from *non-gammaproteobacterial Pseudomonadota*, reflecting its lower prevalence in these taxa. To quantitatively assess whether the distribution of CsrA in our library reflects that of the broader CsrA population, we counted the number of CsrAs in each Phyla in our database and in the library and compared the distribution of CsrA across phyla in both (Fig. 1D). The distribution of CsrA variants in our library closely matches the overall distribution across phyla, with only minor differences. We conclude that our library effectively captures the natural taxonomic diversity of CsrA.

In addition to taxonomic variation, we aimed for our library to be representative of CsrA amino acid sequence diversity. One self-imposed limitation was that to facilitate the commercial synthesis of *csrA* genes we restricted our library to CsrA representatives with fewer than 100 amino acids. However, the size distribution of CsrA proteins in our dataset essentially mirrors that of all bacterial CsrAs, despite the size criterion (Fig. 1E). Further supporting its representativeness, sequence logos comparing all known CsrAs to those in our library reveal similar patterns of conserved residues and relative information, indicating that our dataset captures key functional elements (Fig. 1F). To more quantitatively compare our library to the natural variation in CsrA amino acid sequence we calculated entropy values across CsrA positions. Figure 1G shows that the positional entropy in our library aligns with or is greater than that of all bacterial CsrAs. The increased entropy in our library is likely a consequence of using UniRef50 clustering which ensures variability. Consistent with this, when we randomly sample 550 CsrAs from our database, the positional entropy values are essentially identical to that of all CsrAs (Fig. 1G). As a final test to ensure our library was diverse enough, we performed a pairwise sequence identity comparison for all CsrAs in our library. We find that most CsrAs differ from each other by 30% suggesting that the library captures substantial sequence diversity, as most CsrAs are not too similar to one another (Fig. 1H). The observed normal distribution implies that CsrA sequence diversity in the library is not strongly biased toward highly similar or highly divergent sequences. Together, these analyses confirm that our CsrA library is a robust and representative dataset for functional studies. By encompassing a wide phylogenetic range while preserving key sequence features and diversity, our library provides a powerful framework for systematically probing the functional evolution of CsrA across bacteria.

### Swarm-seq: identification of functional classes of CsrAs

To functionally assess our synthetic library of CsrAs, we developed Swarm-seq, a high-throughput assay that leverages bacterial swarming motility in *B. subtilis* to distinguish functional from non-functional CsrAs. *B. subtilis* normally exhibits robust swarming, but overexpression of its native CsrA (CsrABs) inhibits motility by preventing translation of flagellin called Hag (Yakhnin et al., 2007). CsrABs binds to two GGA-containing stem loops in the 5’-UTR of the *hag* transcript to prevent ribosome binding and translation initiation (Yakhnin et al., 2007) (Fig. 2A), and unlike its Gram-negative counterparts, is antagonized by the protein FliW (Mukherjee et al., 2016, 2013, 2011; Oshiro et al., 2020, 2019). The role of CsrABs is to coordinate *hag* expression with the assembly state of the flagellum, and homeostatically restrict *hag* translation by a negative feedback loop involving FliW and CsrA. Thus, to determine whether *csrAs* from evolutionarily distant bacteria can complement for each other in *B. subtilis,* we engineered a strain where we eliminated the FliW feedback loop and assessed swarming motility upon overproduction of heterologous CsrAs in a Δ*csrA* Δ*fliW* double mutant. This genetic background ensured that swarming motility was directly linked to heterologous CsrAs RNA-binding function. As an initial control, we ensured that this strain was motile when no CsrA was expressed but cells producing CsrABs from an ectopic chromosomal locus upon IPTG induction did not exhibit swarming motility (Fig. 2B). We conclude that overexpression of CsrAs in this background enables the assessment of CsrA inhibitory function.

**Fig. 2:**
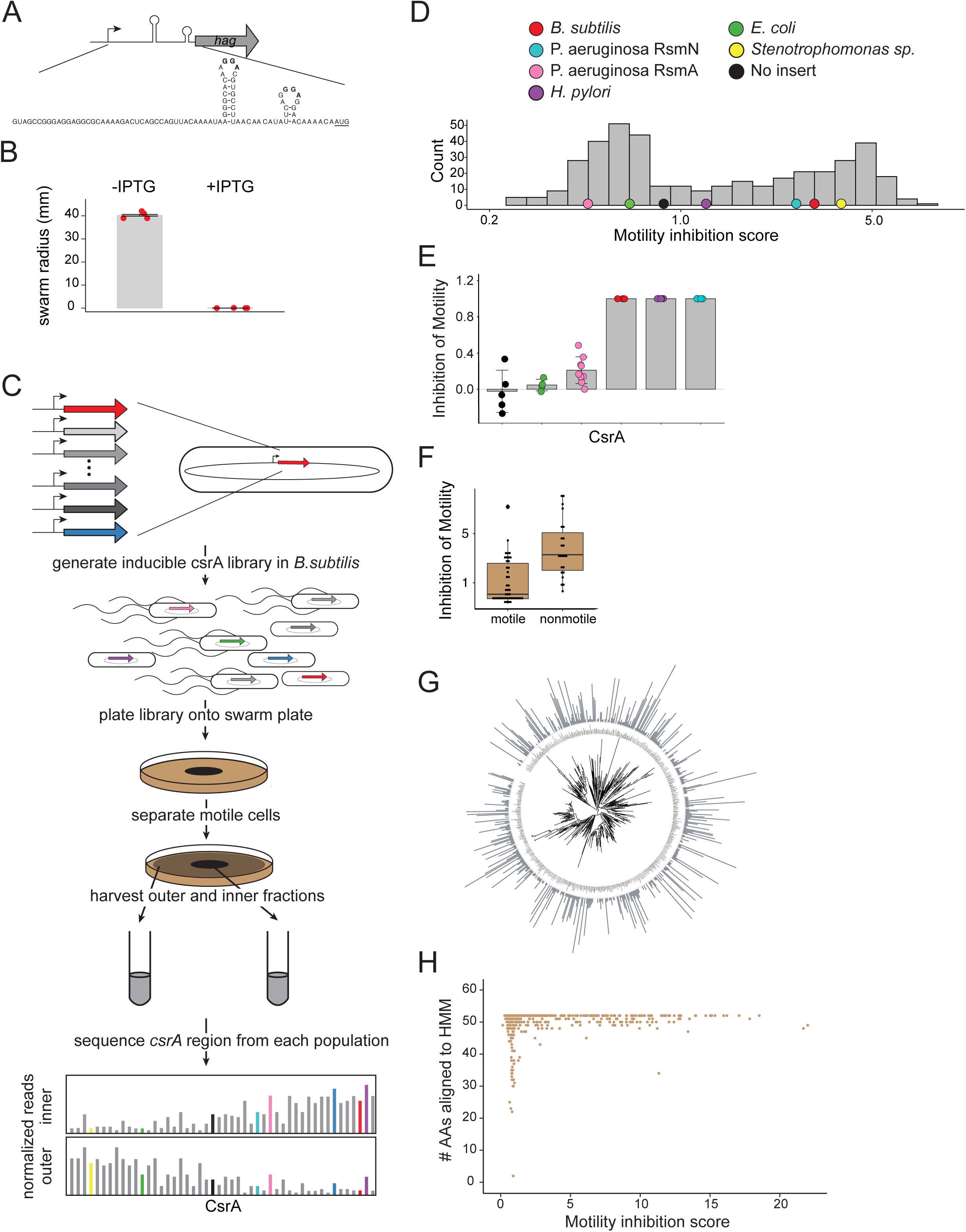
Swarm-seq enables classification of functional and non-functional CsrAs. **(a)** *hag* 5ʹ-UTR sequence and structure. **(b)** Swarm radius quantification after 7 hrs without (-IPTG) and with (+IPTG) overexpression of CsrABs. **(c)** Schematic of the Swarm-seq workflow. **(d)** Histogram of motility scores from Swarm-seq data (Colored circles are representatives colored as in Fig. 1). **(e)** Validation of Swarm-seq data with individual CsrAs. **(f)** Boxplots of Swarm-seq validation examples. **(g)** Motility scores mapped onto tree built with CsrA protein sequences. **(h)** Scatter plot showing number of amino acids that align to the CsrA HMM and the corresponding motility inhibition score for all CsrAs in Swarm-seq data.

We next leveraged CsrA-dependent swarming inhibition to systematically assess all 555 CsrAs in our library (Fig. 2C). First, we generated an IPTG-inducible overexpression library of the *csrA*s. As controls our library included a construct without *csrA* (negative control), and a construct that overexpressed CsrABs (negative control). We grew the library overnight, plated it on the center of a swarm agar plate, allowed the cells to swarm for 7 hrs (the amount of time required for the leading edge of the swarm to reach the outer edge of the plate in the absence of IPTG), then isolated cells from the outer edge of the plate. We purified genomic DNA from the motile outer sub-population and the inner subpopulation of the plate that should contain genetically motile and non-motile cells, amplified the *csrA* locus from both populations, and sequenced the *csrA* locus. We identified reads for over 400 of the CsrAs in the library (Supplementary Fig. 2A). We defined motility inhibition as the number of normalized counts in the inner fraction divided by the number of normalized counts in the motile outer fraction. CsrAs that inhibit motility were expected to be depleted in the motile fraction, while non-functional CsrAs would be evenly distributed or depleted in the non-motile fraction. Figure 2D and Supplementary Fig. 2B show the distribution of motility inhibition scores: as expected, *B. subtilis* expressing its native CsrABs exhibited a high inhibition score, whereas the no-CsrA control had a low inhibition score, suggesting Swarm-seq was accurately separating functional and nonfunctional CsrAs. Surprisingly, the Swarm-seq data indicate that while approximately half of the CsrAs in our library (Class I) showed some level of swarming inhibition, the other half of the CsrA population was unable to inhibit swarming motility (Class II). To validate the Swarm-seq results, we individually isolated strains expressing CsrAs from pathogenic bacteria, including *Escherichia coli*, *Pseudomonas aeruginosa*, and *Helicobacter pylori*, and assessed their effects on swarming in isogenic conditions. Consistent with the Swarm-seq data, we found that CsrABs, RsmN from *P. aeruginosa*, and CsrAHp from *H. pylori* inhibited swarming (Fig. 2E). In contrast, and consistent with the Swarm-seq data, CsrAEc from *E. coli* and RsmA from *P. aeruginosa* did not inhibit swarming (Fig. 2E).

To further test the robustness of Swarm-seq, we grew our library, plated on selective medium, isolated 69 colonies, and individually assessed the swarming behavior of the selected isolates. Of these, 46 isolates were motile when individually assessed and 23 were nonmotile, consistent with the surprising Swarm-seq result that the majority of CsrAs were not able to inhibit motility. We then pooled cells from isolates that were unable to inhibit swarming and separately pooled those from the isolates that successfully inhibited swarming. After growing, amplifying CsrA, and sequencing, we compared the distribution of motility inhibition scores between the two pools. This comparison was consistent with our Swarm-seq data, with the non-swarming isolates showing a significantly higher mean inhibition score (Fig. 2F). These results confirm that Swarm-seq is a sensitive and scalable method for classifying CsrAs based on their ability to regulate motility, providing a functional readout for target recognition and RNA binding across diverse bacterial CsrAs.

To determine whether the ability of CsrA homologs to inhibit motility correlates with phylogeny, we mapped functional and non-functional CsrAs onto a maximum-likelihood phylogenetic tree. Functional and non-functional variants were interspersed throughout the tree, without clustering into distinct clades (Fig. 2G, Supplementary Fig. 3). This pattern suggests that the sequence space compatible with binding and repression of *hag* is broad, but functional activity of CsrA is also sensitive to disruption. Additionally, the high phylogenetic divergence among homologs and the coarse sampling of sequence space likely limit our ability to identify individual amino acid changes that govern functional differences. However, our data clearly show that CsrA homologs that deviate substantially from the CsrA HMM consensus are enriched among sequences that fail to inhibit motility (Fig. 2H).

### Functional CsrAs inhibit expression of *hag*

Swarm-seq was designed to assess if a heterologously expressed CsrA was able to inhibit swarming motility, which is a physiological assay. We hypothesized that functional Class I CsrAs inhibited motility via repression of *hag* translation. To directly test this, we fused the *hag* promoter and 5′ UTR to mNeonGreen, along with a mutant version of the 5′ UTR known to prevent inhibition by CsrABs (Mukherjee et al., 2011) (Fig. 3A). Microscopy confirmed that the reporter functioned as expected, with green fluorescence observed in the absence of CsrABs or when IPTG was not added, while fluorescence signal was diminished upon IPTG induction of CsrABs (Fig. 3B). We then tested different strains, including one without CsrA, one expressing CsrABs, and strains expressing CsrAEc, RsmA, RsmN, CsrASm (*S. maltophilia* CsrA), and CsrAHp. Consistent with the Swarm-seq data, we found that CsrAEc, RsmA, and the no-CsrA control were unable to inhibit wildtype *hag-mNeonGreen*, while CsrABs, RsmN, CsrASm, and CsrAHp successfully inhibited *hag* reporter expression. Notably, while most CsrAs lost the ability to inhibit the mutant 5’ UTR, CsrASm and CsrAHp retained inhibition (Fig. 3C). These results suggest that while some CsrAs share the ability to repress *hag* translation, others exhibit distinct regulatory specificities, with *S. maltophilia* and *H. pylori* CsrAs potentially recognizing additional sequence or structural features beyond those required by *B. subtilis* CsrA or their binding not being dependent on the same determinants as CsrABs.

**Fig. 3:**
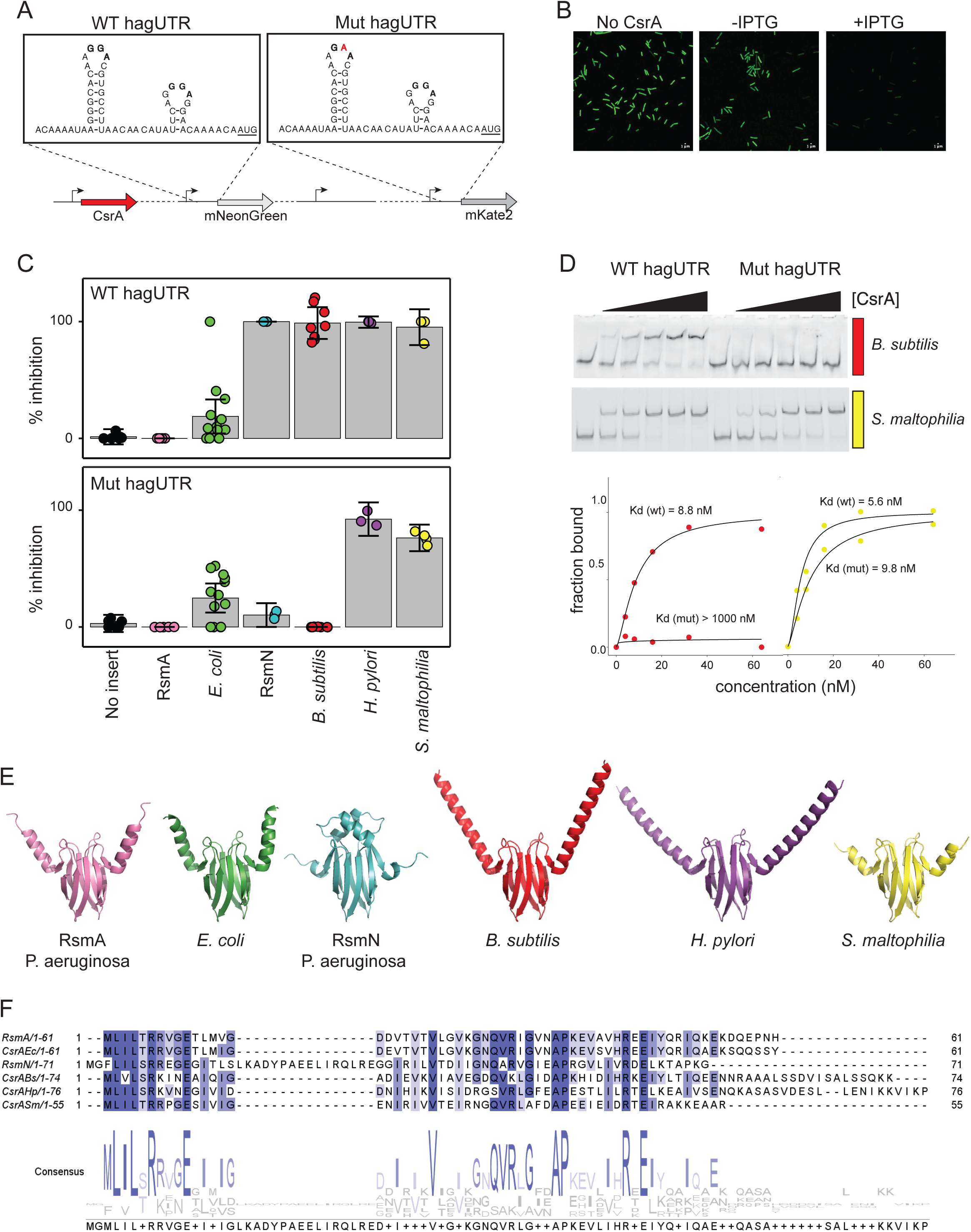
Functional CsrAs directly inhibit expression by binding the *hag* 5ʹ-UTR. **(a)** Sequence and secondary structure of wild-type (left) and mutant (right) hag 5ʹ-UTR. The G to A substitution that eliminates binding by CsrABs is highlighted in ref. P_hag_ sequence from −144 to +10 were fused to genes coding for mNeonGreen and mKate2. **(b)** Representative green fluorescence micrographs of reporter strain without (-IPTG) and with (+IPTG) overexpression of CsrABs. **(c)** Inhibition of expression of representative CsrAs (colors as in Figs. 1 and 2). Expression was normalized to no CsrA control done in same 96-well plate. Inhibition is 1-fraction of expression of no CsrA. **(d)** Electrophoretic mobility shift assays and binding quantification of CsrABs and CsrASm on RNA fragments with WT sequence (left) from −60 to +15 or the GGA->GAA mutant sequence (right) from −60 to +15. **(e)** Alphafold3 models of representative CsrAs. **(f)** Amino acid sequence alignment of proteins shown in (e).

To further investigate the unexpected specificity observed in our reporter assays, we purified CsrABs and CsrASm and assessed their RNA-binding capabilities using an electrophoretic mobility shift assay (EMSA). When incubated with wildtype (WT) *hag* 5′UTR RNA, CsrABs exhibited strong binding but showed little to no interaction with the mutant *hag* 5′UTR, indicating that it can effectively discriminate between the two target RNA sequences (Fig. 3D). In contrast, CsrASm bound both the WT and mutant *hag* 5′UTRs with comparable affinity, with less than a two-fold difference (Fig. 3D). These results suggest that while CsrABs exhibits sequence-specific recognition of the *hag* 5′UTR, CsrASm has a broader binding profile that may allow it to regulate a more diverse set of targets.

To investigate whether structural features might explain differences in CsrA function, we predicted the structures of several CsrA homologs — CsrABs, CsrAEc, RsmA, RsmN, CsrAHp, and CsrASm — using AlphaFold3 (Abramson et al., 2024) (Fig. 3E). The most apparent structural difference among these homologs is the length of the C-terminal helices, which serve as binding sites for regulatory proteins (such as FliW) in some species, including *B. subtilis*. Another prominent feature is unique to RsmN: an enlarged loop containing an additional α-helix near the RNA-binding site (visible in the top-middle portion of the structure in Fig. 3E). However, this feature does not correlate with functional differences, as RsmN behaves similarly to CsrABs in our assays. In addition, aligning the primary sequence of the CsrAs shown in Fig. 3E did not exhibit a clear pattern that enabled prediction of function. For example, RsmN which was only 24.53% identical complemented the lack of CsrABs, while RsmA which was 41% identical did not (Fig. 3F). The overall inability to predict function from sequence or structural analysis highlights the importance of high-throughput functional assays like Swarm-seq.

### *B. subtilis* as a chassis for CsrAs from challenging-to-study bacteria

Our library includes several CsrAs from industrially and environmentally significant bacteria spanning the bacterial tree (Fig. 4A). Among these, *Congregibacter litoralis* CsrA is particularly noteworthy due to its role in aerobic anoxygenic phototrophy, a process that influences marine carbon cycling (Spring et al., 2009); however, studying its regulation in its native host is complicated by its reliance on specialized metabolic pathways. Similarly, *Blochmannia floridanus*, an obligate endosymbiont of carpenter ants, presents unique challenges as it lacks many genes required for independent survival, making genetic manipulation difficult (Feldhaar et al., 2007). As another example, *Pseudomonas chlororaphis* is an agriculturally important biocontrol bacterium known for producing antifungal compounds, yet its complex regulatory networks and secondary metabolite production pathways complicate efforts to dissect CsrA function (Calderón et al., 2015). Finally, *Nitrospira defluvii*, a key organism in wastewater treatment due to its role in complete ammonia oxidation (comammox), grows slowly, making functional studies in its native context particularly challenging (Latocheski et al., 2022). Using our heterologous system, we demonstrated that CsrAs from these challenging-to-study bacteria cluster within the functional region of our motility score histogram, indicating their ability to inhibit motility in *B. subtilis* (Fig. 4B). Structural predictions and sequence alignments for these functional CsrA proteins revealed highly similar overall folds and sequence identity, but the *N. defluvii* protein was predicted to have an extra loop in the RNA binding interface that is surprising considering its functionality (Fig. 4C, D). This further highlights the inability to predict CsrA function from predicted structure.

**Fig. 4:**
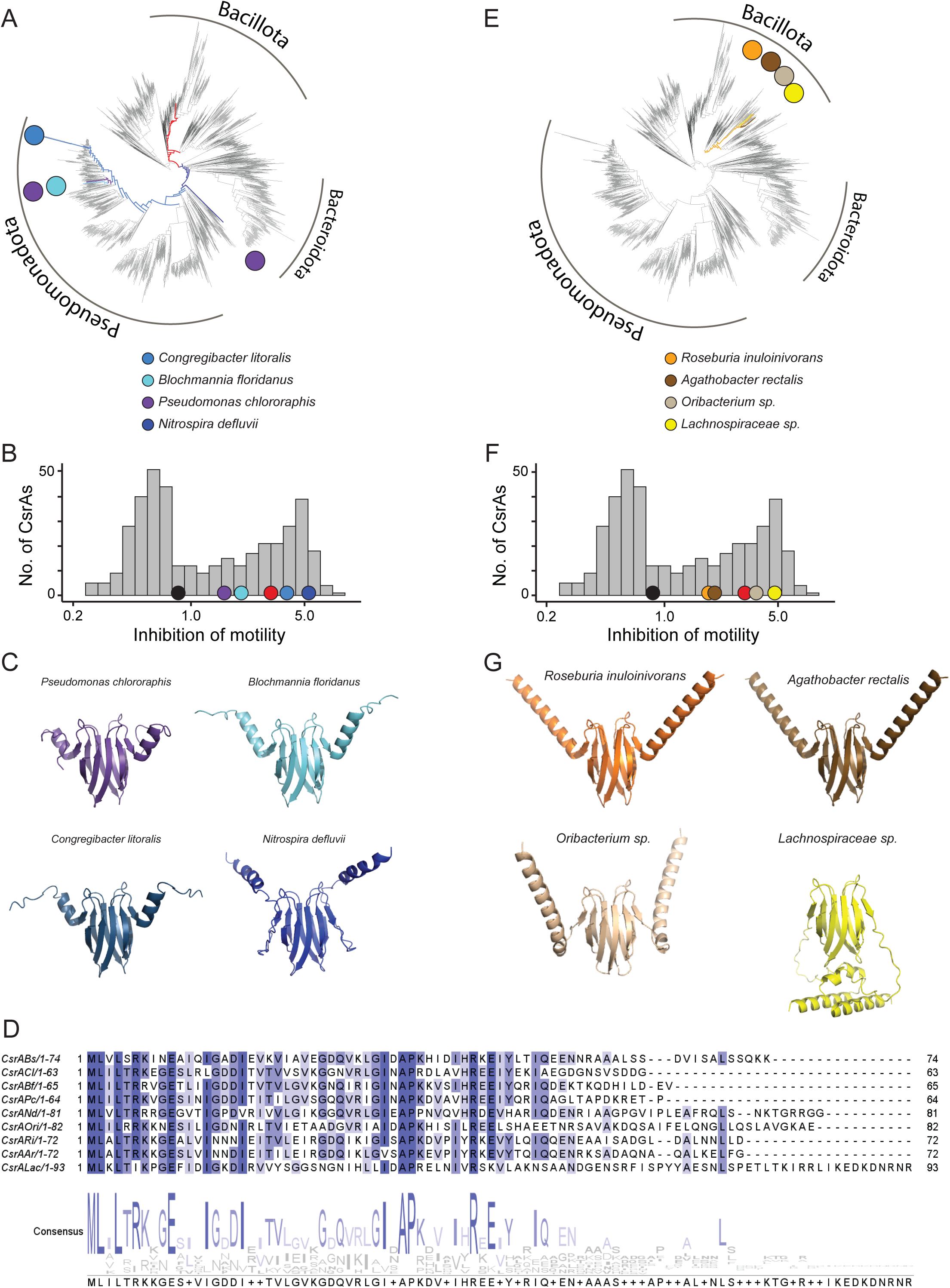
Swarm-seq enables the study of industrially relevant bacterial CsrAs. **(a)** Phylogenetic tree of species containing CsrA as in Fig. 1a. Colored circles and branches highlight genera containing industrially relevant CsrAs that are functional in the Swarm-seq dataset. **(b)** Motility inhibition scores for relevant CsrAs are shown in histogram of scores. Circles show position of representative CsrAs within histogram. **(c)** Structural models of highlighted CsrAs generated with AlphaFold3 using dimer inputs. **(d)** amino acid sequence alignments of select CsrA proteins. **(e)** Phylogenetic tree of species containing CsrA as in Fig. 1a. Colored circles and branches highlight genera containing microbiome CsrAs that are functional in the Swarm-seq dataset. **(f)** Motility inhibition scores for relevant CsrAs are shown in histogram of scores. Circles show position of representative CsrAs within histogram. Colors as in (a). **(g)** Structural models of highlighted CsrAs generated with AlphaFold3 using dimer inputs.

Similarly, our library includes several CsrAs from microbiome-associated bacteria, including *Roseburia inulinivorans*, *Agathobacter rectalis*, an *Oribacterium* species, and another uncharacterized *Lachnospiraceae* species (Fig. 4E). *Roseburia inulinivorans* and *Agathobacter rectalis* are key butyrate-producing bacteria in the human gut, contributing to intestinal health and immune regulation (Singh et al., 2023; Vital et al., 2017). *Oribacterium* species are members of the oral microbiome, with potential roles in maintaining microbial balance and interacting with host tissues (Sizova et al., 2014). *Lachnospiraceae* species are abundant in the gut and associated with fiber fermentation and metabolic health, but many remain poorly characterized (Vacca et al., 2020). Using Swarm-seq, we demonstrated that these CsrAs cluster within the functional region of our motility score histogram, indicating their ability to inhibit motility in *B. subtilis* (Fig. 4F, G). By characterizing these diverse CsrAs in a genetically tractable system, we provide insights into their regulatory potential that would be difficult to obtain in their native hosts.

## DISCUSSION

Our findings fundamentally reshape our understanding of CsrA-mediated post-transcriptional regulation across bacterial species. The functional divergence we observed among CsrA homologs challenges the long-standing assumption that these highly conserved RNA-binding proteins are functionally interchangeable between species. The inability of well-characterized CsrA variants from *E. coli* and *P. aeruginosa* to substitute for native *B. subtilis* CsrA in regulating flagellar motility was particularly surprising given the high sequence conservation (>40% identity) among these proteins. This observation suggests that subtle differences in protein structure or RNA-binding preferences have evolved to accommodate species-specific regulatory needs. Such functional specialization likely reflects the distinct ecological niches and physiological demands of different bacterial species, underscoring the remarkable adaptability of bacterial regulatory systems despite their seemingly conserved components. Furthermore, our phylogenetic analysis revealed that functional and non-functional CsrAs are interspersed throughout the evolutionary tree rather than clustering in specific clades, suggesting that functional divergence has arisen multiple times independently. Although our synthetic CsrA library is too phylogenetically coarse to pinpoint the specific sequence changes in CsrA or its RNA targets that underlie this divergence, our findings highlight the remarkable plasticity of the CsrA–RNA interaction. Future efforts will require finer-resolution phylogenetic sampling and targeted mutational analyses to identify the molecular determinants that drive these functional transitions.

We demonstrated striking differences in the ability of various CsrA homologs to inhibit translation. A consensus sequence for *E. coli* CsrA binding sites was determined as RUACARGGAUGU using systematic evolution of ligands by exponential enrichment (SELEX)(Dubey et al., 2005) Similarly, a SELEX approach combined with deep sequencing of RNAs identified CANGGAYG with the GGA motif typically found in a hexaloop as consensus for RsmA and RsmF(Schulmeyer et al., 2016b) Nonetheless, we show that while *E. coli* CsrA and *P. aeruginosa* RsmA failed to inhibit wildtype *hag* expression, *P. aeruginosa* RsmN, *S. maltophilia* CsrA, and *H. pylori* CsrA successfully repressed translation. Most notably, *S. maltophilia* and *H. pylori* CsrAs retained inhibitory function even with the GGA → GAA mutant UTR that is blind to regulation by *B. subtilis* CsrA. Our EMSA results confirmed these functional differences. These findings suggest that some CsrA homologs have evolved highly specific RNA recognition patterns, while others maintain broader binding profiles enabling regulation of more diverse targets.

Swarm-seq represents a significant methodological advancement that overcomes longstanding technical barriers in studying post-transcriptional regulation across diverse bacterial species. By providing a standardized genetic background and a quantifiable phenotypic readout, our approach isolates the functional contribution of each CsrA variant with unprecedented precision. This enables comparative analyses that were previously impossible, particularly for bacteria that are genetically intractable or difficult to culture in laboratory settings. The scalability of Swarm-seq further enhances its utility, allowing for high-throughput screening of CsrA homologs from hundreds of species simultaneously with the potential to scale to many thousands of species. Our ability to characterize CsrA function in environmentally and medically relevant bacteria — from marine carbon cycling specialists to human gut microbiome members — demonstrates the method’s versatility. These insights into previously inaccessible regulatory networks may have significant implications for fields ranging from environmental microbiology to human health, potentially informing strategies for manipulating bacterial behavior in various contexts.

The implications of our findings extend from fundamental evolutionary biology to practical applications in synthetic biology and medicine. The ability to predict which CsrA homologs will function in heterologous hosts could facilitate the design of synthetic regulatory circuits with enhanced portability across bacterial species. This knowledge could accelerate the development of engineered bacteria for applications ranging from bioremediation to therapeutic delivery, by enabling more reliable transfer of regulatory components between chassis organisms. Furthermore, the species-specific functionality we observed suggests that therapeutic strategies targeting CsrA-mediated regulation may need to be tailored to individual pathogens rather than applying insights from model organisms like *E. coli*.

The functional conservation we observed among CsrA homologs from evolutionarily distant species such as the environmentally significant bacteria and microbiome-associated species highlighted in Figure 4 is equally intriguing. CsrALac had a higher motility inhibition score than CsrARi despite CsrALac having a lower sequence identity to CsrABs (31.08%) when compared to CsrARi (48.65%) highlighting the need for a functional assay such as Swarm-seq. This functional conservation across species that are evolutionarily distant and technically challenging to study in their native contexts highlights the power of Swarm-seq as a platform for investigating post-transcriptional regulation in difficult-to-manipulate bacteria. Identifying functional CsrA-RNA interaction principles using Swarm-seq could provide crucial insights into the fundamental principles governing RNA-protein interactions in the native bacterium and potentially inform the design of synthetic RNA regulatory systems with predictable behaviors in different species.

## MATERIALS AND METHODS

### Strains and growth conditions

*B. subtilis* and *E. coli* strains were grown in lysogeny broth (LB) (10 g tryptone, 5 g yeast extract, 5 g NaCl per liter) broth or on LB plates fortified with 1.5% Bacto agar at 37 °C for most experiments. For reporter assays, *B. subtilis* strains were grown in 1% tryptone. The following antibiotic concentrations were used when necessary: ampicillin 100 μg/ml (*amp*), kanamycin 5 μg/ml (*kan*), chloramphenicol 5 μg/ml (*cm*), spectinomycin 100 μg/ml (*spec*), and tetracycline 10 μg/ml (*tet*).

### Strain construction

#### Construction of ycgO::Phag(wt)-mNeonGreen–tet

To generate a translational reporter using the wild-type *hag* promoter and 5′ UTR, we amplified the region from −144 to +15 relative to the *hag* start codon from *B. subtilis* NCIB3610 genomic DNA using primers JW13 and JW14. The mNeonGreen gene was amplified using primers JW15 and JW16. These two fragments were joined via Gibson assembly, and the product was further amplified using primers JW13 and JW16 to yield a 918 bp insert. The plasmid backbone was generated by amplifying pKM086 (a gift from Dan Kearns) with primers JW12 and JW17, producing a 6704 bp linear fragment. The insert and backbone were assembled by Gibson assembly and transformed into chemically competent *E. coli* DH5α. The correct clone was confirmed by whole plasmid sequencing (Plasmidsaurus). The resulting plasmid was transformed into *B. subtilis* strain DK9452 and plated on LB + 10 μg/mL tetracycline, yielding strain SBs32.

#### Construction of yuxG::Phag(GGA->GAA)-mKate2–cm

To generate a second translational reporter using the GGA->GAA variant of the *hag* 5′ UTR, the region from −144 to +15 was amplified from DS6530 genomic DNA using primers JW23 and JW24. The mKate2 gene was amplified from plasmid pNUT029 using primers JW21 and JW22. Gibson assembly was performed on the two PCR products, and the resulting fragment was amplified with JW21 and JW24. Plasmid pWX144 (gift from Dan Kearns) was amplified using primers JW20 and JW25 to provide the backbone. The insert and backbone were assembled by Gibson assembly and transformed into chemically competent *E. coli* DH5α. Following sequence confirmation (Plasmidsaurus), the plasmid was transformed into strain SBs32 and selected on LB + 5 μg/mL chloramphenicol, generating strain SBs36.

#### IPTG-inducible constructs for expression in B. subtilis

To generate the inducible *amyE*::*P_hyspank_-csrA spec* constructs described below, pLC109 (Mukherjee et. al.2011) was used as a backbone for Gibson assemply with the PCR products described below. The resulting plasmids were transformed into DK9452 for swarming assays or SBs36 for reporter assays and selected for using 100 µg/ml spectinomycin.

To generate amyE::Physpank-csrASm spec, *csrA* was amplified from the oligo pool using primers JW224 and JW225. Transformation into DK945 resulted in strain SBs91. Transformation in SBs36 resulted in strain SBs92.

To generate amyE::Physpank-rsmA spec, *rsmA* was amplified from the oligo pool using primers JW155 and JW156. Transformation into DK945 resulted in strain SBs50. Transformation into SBs36 resulted in strain SBs60.

To generate amyE::Physpank-rsmN spec, *rsmN* was amplified from the oligo pool using primers JW157 and JW158. Transformation into DK945 resulted in strain SBs51. Transformation into SBs36 resulted in strain SBs61.

To generate amyE::Physpank-csrAEc spec, csrAEc was amplified from the oligo pool using primers JW159 and JW160. Transformation into DK9452 resulted in strain SBs54. Transformation into SBs36 resulted in strain SBs65.

To generate amyE::Physpank-csrAHp, *csrA*Hp was amplified from the oligo pool using primers Transformation into DK945 resulted in strain SBs83. Transformation into SBs36 resulted in strain SBs45.

#### csrA overexpression in E. coli

To generate the inducible pET28b-*csrA kan* constructs described below, pET28b amplified with EY13 and EY14 was used as a backbone for Gibson assemply with the PCR products described below. The resulting plasmids were transformed into BL21(DE3) and selected for using 50 µg/ml kanamycin.

To generate pET28b-*csrA*Bs, *csrABs* was amplified from the oligo pool using primers EY11 and EY12.

To generate pET28b-*csrA*Sm, *csrASm* was amplified from the oligo pool using primers JW224 and JW225.

### Local Genome Database and Representative Tree

We constructed a local genome database by downloading representative genomes from the ProGenomes v3 (Fullam et al., 2023). To ensure high-quality annotations, we retained only genomes with available NCBI annotations, yielding a final dataset of 26,374 annotated genomes. For phylogenetic context (Fig. 1B), we downloaded the bac120 r214 species tree in Newick format from the Genome Taxonomy Database (GTDB;(Parks et al., 2022)). Each genome in our local database was matched to its corresponding leaf in the GTDB tree, and GTDB taxonomic assignments were used for classification at the phylum, class, order, family, genus, and species levels. To generate a reduced genus-level tree, we randomly selected one genome per genus and pruned all other leaves using the Python library ETE3 v3.1.2(Huerta-Cepas et al., 2016).

### Identification of CsrA Homologs

To identify putative CsrA homologs within our local genome database, we used hmmsearch v3.3.2(Eddy, 2011) to scan all proteomes against the PF02599 CsrA HMM profile. Protein hits with an E-value below 1e-5 were retained for downstream analyses. These identified sequences were then realigned to the HMM profile using hmmalign, and unaligned regions were trimmed from the alignment ends. Shannon entropy for each alignment position was calculated using the formula: H=−∑pilog 2(pi)H = -\sum p_i \log_2(p_i)H=−∑pilog2(pi) where pip_ipi is the frequency of amino acid iii at a given alignment position. Pairwise distance matrices were generated using Biopython’s DistanceCalculator module (Cock et al., 2009) with the ‘identity’ metric.

### Phylogenetic Analyses

Species trees were inferred following a pipeline similar to Parks et al. (2022). Python code for constructing the full bac120 tree from proteomes and HMM profiles is available at https://github.com/JonWinkelman/bac120_tree_inference. For the CsrA gene tree, we aligned all curated CsrA sequences to the PF02599 HMM, trimmed unaligned terminal residues, and removed alignment columns where more than 90% of sequences contained gaps. Phylogenetic trees were inferred using FastTree v2.1.11(Price et al., 2010) with the LG amino acid substitution model(Le and Gascuel, 2008). Final trees were visualized and annotated using iTOL v6(Letunic and Bork, 2007).

### Selection of *csrA*s to be synthesized

To construct a diverse and non-redundant library of CsrA-like sequences, we identified UniRef50 clusters(Suzek et al., 2015) where the representative sequence was annotated as a “Translational regulator CsrA,” yielding over 1,500 clusters. From these, we selected the representative sequence from 550 distinct clusters to include in our curated CsrA library. Sequences were aligned to the PF02599 CsrA hidden Markov model (HMM) profile from the Pfam database(Mistry et al., 2021) using hmmalign v3.3.2. Selected *csrA* genes were codon optimized for expression in *B. subtilis*. Constant sequences were added to the 5ʹ (5ʹ-CTAAAGTCACGGAGGTGTCAGTCAC-3ʹ) and 3ʹ (5ʹ-AAGTCACAAAGTCAGAAATCGTGGC-3ʹ) ends of the coding sequences to amplify and clone the *csrA*s. The pool of 555 sequences were purchased from IDT.

### IPTG-inducible *csrA* library construction

Sequences from the csrA oligo pool were amplified using primers JW76 and JW201. Plasmid pLC109 a Physpank promoter and a spectinomycin resistance gene inserted into amyE locus was digested with NheI and SalI. NheI- and SalI-digested Oligo pool PCR products and plasmid backbone PCR products ligated and transformed into MegaX DH10B T1^R^ electrocompetent cells (Invitrogen C640003). After a 1 hr recovery, serial dilution plating onto LB plates containing 100 µg/ml carbenicillin was performed to approximate the number of transformants. The remaining transformation was grown overnight in liquid LB with 100 µg/ml carbenicillin. Plasmid was purified from the overnight culture with a Qiagen plasmid miniprep kit following the manufacturer’s instructions. We next introduced the plasmid into *B. subtilis* strain DK9452 using natural competence. All transformants were selected for on LB agar plates with 100 µg/ml spectinomycin then harvested from the plates.

### Swarm expansion assay

Cells were grown to mid-log phase at 37°C in LB broth and resuspended to 10 OD_600_ in PBS buffer (137 mM NaCl, 2.7 mM KCl, 10 mM Na2HPO4, and 2 mM KH2PO4) containing 0.5% India ink ().LB containing 0.7% Bacto agar was prepared the day before the experiment and 25 ml was added to each petri dish. For all experiments, swarm plates were made with and without 1 mM IPTG. On the day of the experiment, the plates were dried for 5 min in a laminar flow hood before being centrally inoculated with 10 µl of the cell suspension, and dried for another 5 min. For all experiments, a single cell suspension was added to a plate with IPTG and a plate without IPTG. Plates were incubated at 37°C in an incubator with humidity maintained over 40% for 7 hrs before measuring the radius of the swarm edge. The India ink demarcated the origin of the colony and the swarm radius was measured relative to the origin. To define motility inhibition, the swarm radius for each cell suspension was calculated on the +IPTG plate and the -IPTG plate. Motility inhibition was then defined as:

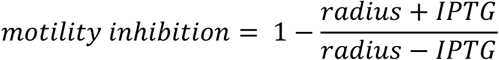

### Swarm-seq procedure

Cells containing the Physpank-*csrA* library were grown to mid-log phase at 37°C in LB broth and resuspended to 10 OD_600_ in PBS buffer (137 mM NaCl, 2.7 mM KCl, 10 mM Na2HPO4, and 2 mM KH2PO4). Plates were prepared fresh as described above. Cells were incubated for 4 hrs, resulting in a swarm leading-edge that reached within 10 cm of the edge of the plate. Cells from the leading edge were harvested with a sterile cotton swab and inoculated into LB broth. Cultures were grown O/N at 37° with shaking. Genomic DNA was harvested from 1 ml of the O/N culture and 40 ng of genomic DNA was used as template for a 15 cycle PCR with primers JW194 and JW207. The resulting PCR product was amplified in a 15 cycle PCR using primers JW197 and either JW196, JW210, JW211, JW212, or JW213. These primers contain 3 nt barcodes that allowed us to pool the resulting PCRs and amplify the pool with primers JW198 and JW199 then assign the resulting sequences to a specific sample. This final PCR was sequenced on an Illumina Hiseq with 2 x 150 by Seqcoast genomics.

### Swarm-seq analysis

For each fastq file, raw fastq reads were searched for the constant upstream sequence 5ʹ-CTAAAGTCACGGAGGTGTCAGTCAC-3ʹ. Reads having a perfect match were filtered and the 25 nt sequence downstream of the constant sequence was then used to assign the *csrA* in each read. Each *csrA* was then counted and normalized as a fraction of total reads in the library. For each *csrA*, the normalized count in the file from the outer fraction was then divided by the normalized count from the file for the inner fraction. A motility inhibition score was then defined as:

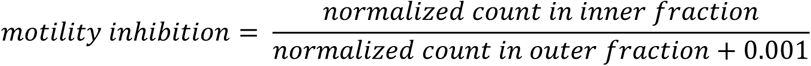

### Fluorescent reporter assay

To measure the effect of CsrA on expression from Phag with a wildtype hag 5ʹUTR and the GGA->GAA mutant hag 5’UTR we grew strains SBs36 in 200 µl of 1% Tryptone broth without the addition of IPTG in a 96-well plate in plate reader for 16 hrs with filters for mNeonGreen and mKate2. Wells were then diluted 10 µl into 190 µl of 1% Tryptone broth with 1 mM IPTG and grown with orbital shaking for 6 hrs at 37℃ while reading the OD and fluorescence every 15 min. Fluorescence/OD readings at mid-log phase were calculated and inhibition scores were defined as:

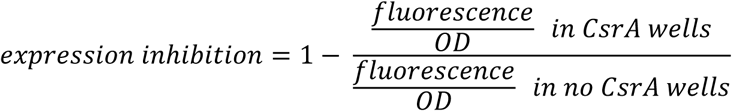

### Purification of His-tagged CsrAs

BL21(DE3) *E. coli* cells containing were grown at 37°C in LB broth and 50 µg ml^−1^ of kanamycin to an OD_600_ of 0.6, at which time 1 mM IPTG was added to the culture and incubation was continued for 3 h. Cells were harvested by centrifugation and the cell pellet was suspended in 20 lysis buffer containing 50 mM Na_2_HPO_4_, 500 mM NaCl, 20 mM imidazole, and 0.1%Tween 20. Cell lysate was prepared by adding 2 ml Bug Buster supplemented with 2 µl Benzonase nuclease, followed by centrifugation at 14,000 rpm for 30 min at 4°C. The resulting supernatant was incubated with 1 ml of Qiagen NiNTA slurry that had been pre-equilibrated with 50 mM Na_2_HPO_4_, 500 mM NaCl, 20 mM imidazole, and 0.1%Tween20 then applied to a 10 ml gravity column. The column was washed with 10 ml 50 mM Na_2_HPO_4_, 500 mM NaCl, 75 mM imidazole, and 0.1%Tween20. His-CsrA was eluted with 50 mM Na_2_HPO_4_, 500 mM NaCl, 275 mM imidazole, and 0.1%Tween20. Elutions were separated by SDS-PAGE and stained with Coomassie Brilliant Blue to verify purification of the His-CsrA fusion. Fractions containing pure His-CsrA were combined and dialysed against 20 mM Tris-HCl, pH 8.0, 200 mM NaCl, 10% Glycerol and 1 mM EDTA and stored at −80°C.

### Electrophoretic Mobility Shift Assay

To generate a template for in vitro transcription of WT hagUTR RNA, oligo EY86 was amplified with JW100 and JW101, To generate templates for in vitro transcription of mutant hagUTR EY87 was amplified with JW100 and JW101. After purification using a Qiagen PCR purification kit, RNA was transcribed in vitro using an NEB HiScribe® T7 High Yield RNA Synthesis Kit following the manufacturer’s instructions. DNaseI treated RNA was then phenol extracted, and ethanol precipitated before being resuspended in RNAse free water. To label the RNAs, Cy5 5ʹ-end-labeled oligo JW71 was annealed to 100 nM of wt or sow3 RNA in 10 mM Tris-Cl 50 mM NaCl by heating to 90℃ then rapidly cooling to 25℃ in a thermocycler. Serial dilutions of CsrABs or CsrASm were bound in 10 µl reactions with 0.4 nM RNA in 25 mM Tris-Cl pH 8.0, 150 mM NaCl, 3 mM MgCl_2_, and 0,01% tween20. After incubating for 5 min at 25℃, 2 µl of EZ-vision One DNA dye was added to the binding reactions and the whole reaction was run on a 15% acrylamide 0.5X TAE gel electrophoresed in 0.5X TAE. Gels were then visualized using a Typhoon imager and quantified using ImageQuant.

### Structural Prediction

Multimer structure prediction of CsrA dimers was performed using Alphafold3 (Abramson et al., 2024) and the top ranked model was visualized using the PyMOL Molecular Graphics System, Version 2.5.0 (Schrödinger, LLC).

## ACKNOWLEDGEMENTS

We thank Dan Kearns for the generous gift of *B. subtilis* strains and plasmids, and thoughtful discussions. Research reported in this publication was supported by the National Institute of General Medical Sciences of the National Institutes of Health (NIH) under Grants R35GM150803 and R00GM129424 and the Searle Scholars Program Grant SSP-2022-104 to S.M. The content of this study is solely the responsibility of the authors and does not necessarily represent the official views of the funding agencies. The funders had no role in study design, data collection and analysis, decision to publish, or preparation of the manuscript.

## SUPPLEMENTARY INFORMATION

**Supplementary Fig. 1:**
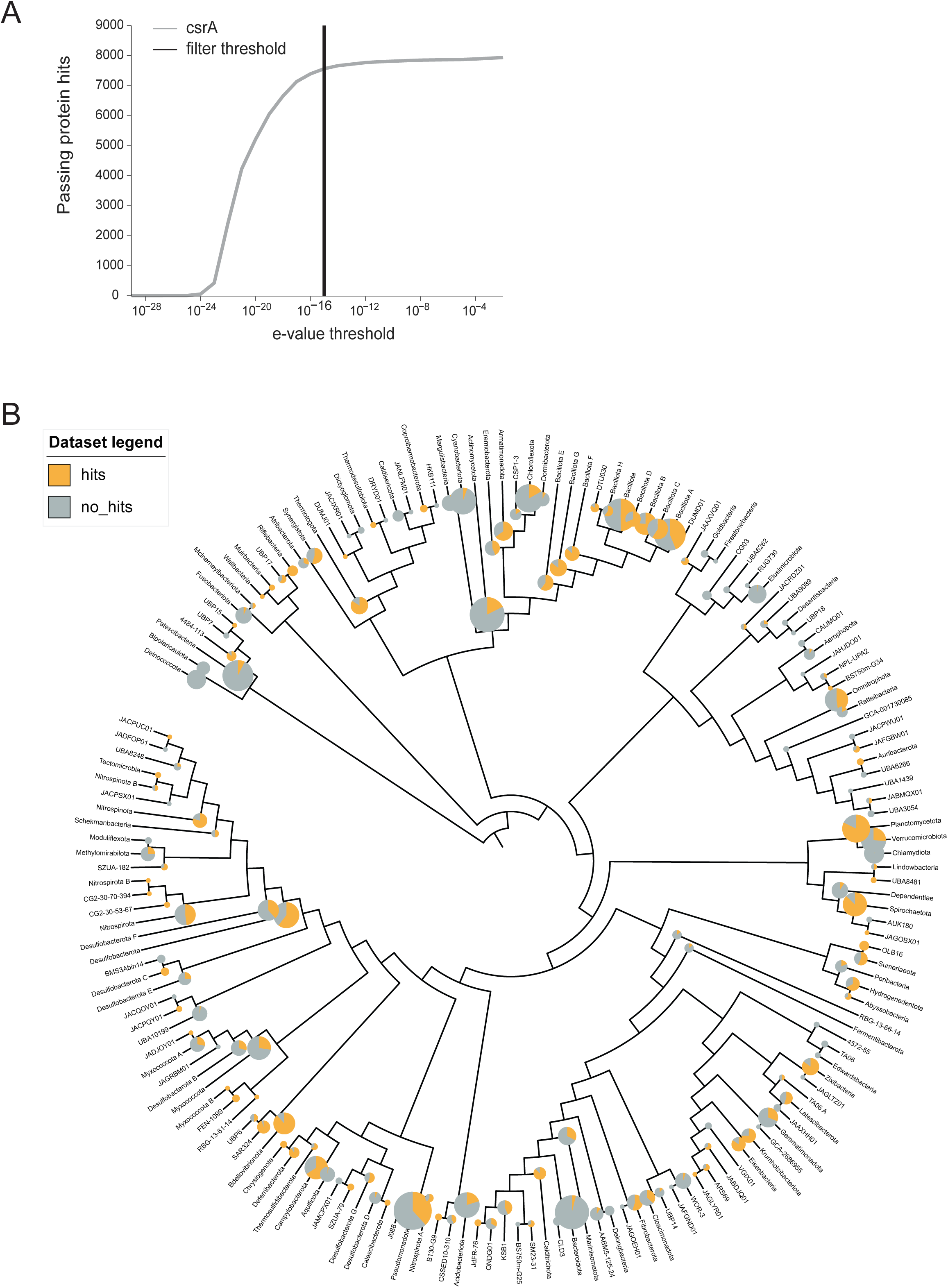
Synthetic *csrA* library is representative. **(a)** Number of CsrAs identified as a function of HMM E-value filter. Vertical line represents filter chosen for this work. **(b)** Phylogenetic tree showing the fraction of species in each phylum that contains CsrA is represented by the pie chart. The size of the pie chart represents the number of species in each phylum.

**Supplementary Fig. 2.**
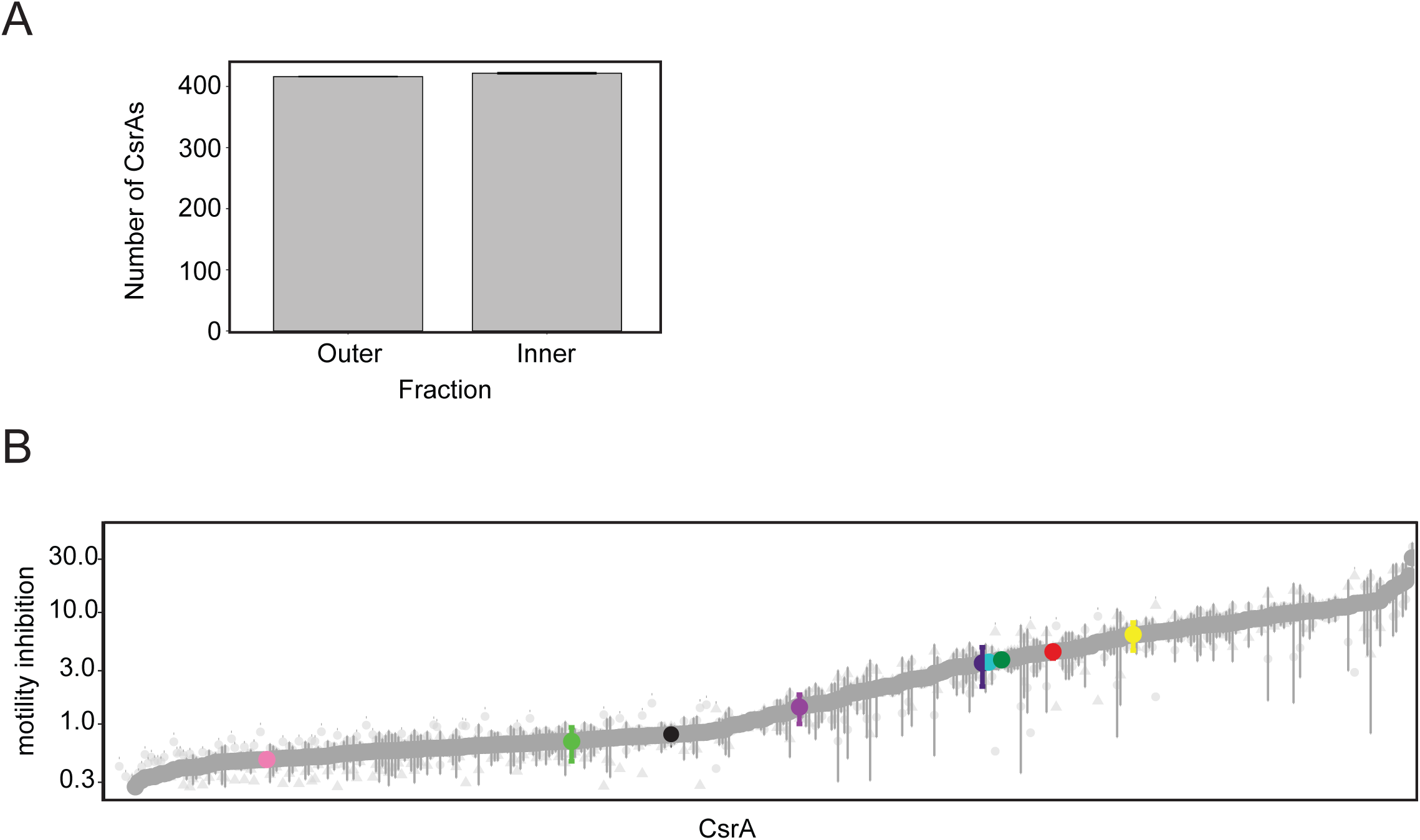
Swarm-n-seq separates CsrAs that inhibit motility from those that do not. **(a)** The number of CsrAs identified by sequencing the library inner and outer fractions. **(b)** Motility inhibition score for each CsrA is plotted from lowest mean motility inhibition score on the left to highest mean motility inhibition score on the right. The mean is a dark gray circle. Data points from individual biological replicates are light gray circles, triangles, or squares depending on the replicate. Bars represent SEM, Colors as in Fig. 1.

**Supplementary Fig. 3.**
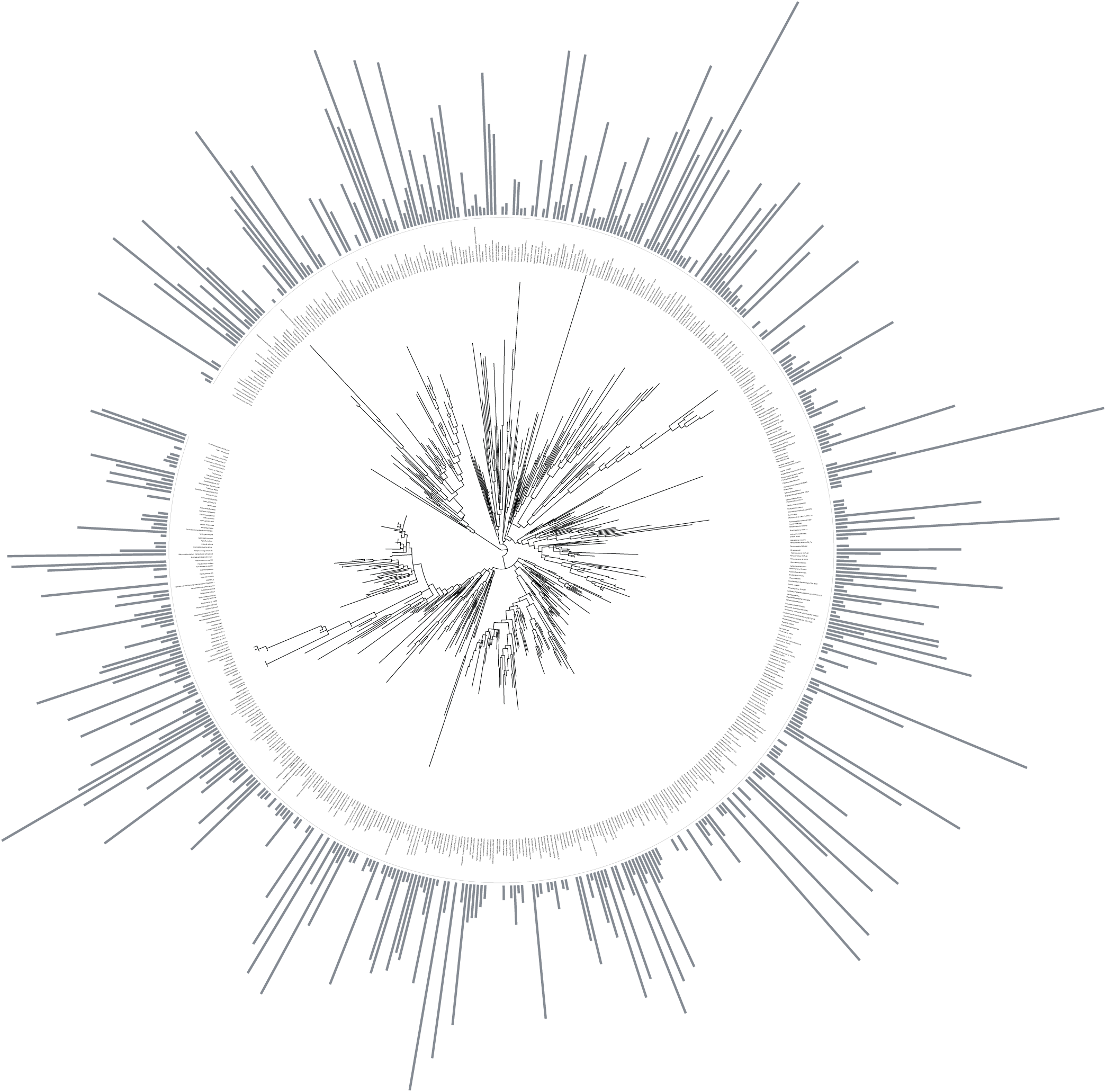
Functional scores do not segregate based on phylogeny. Protein tree of all CsrAs in the library. Length of gray lines represents the motility inhibition score.

